# HTS-Oracle: A Retrainable AI Platform for High-Confidence Hit Identification Across Difficult-to-Drug Targets

**DOI:** 10.1101/2025.07.21.666047

**Authors:** Hossam Nada, Laura Calvo-Barreiro, Sungwoo Cho, Saurabh Upadhyay, Nicholas A Meanwell, Moustafa T. Gabr

## Abstract

Despite rapid advances in computational drug discovery, high-throughput screening (HTS) remains the primary method for identifying initial hits, particularly for targets with limited tractability to small molecules. Yet conventional HTS campaigns are costly and inefficient, often yielding hit rates below 2% and discarding valuable negative data. Here we present HTS-Oracle, a retrainable, deep learning–based platform that integrates transformer-derived molecular embeddings (ChemBERTa) with classical cheminformatics features in a multi-modal ensemble framework for hit prediction. We applied HTS-Oracle to the immune co-stimulatory receptor CD28, a prototypical difficult-to-drug target, and prioritized 345 candidates from a chemically diverse library of 1,120 small molecules. Experimental screening via temperature-related intensity change (TRIC) identified 29 hits (8.4% hit rate), representing an eightfold improvement over conventional methods such as surface plasmon resonance (SPR), TRIC, and affinity selection mass spectrometry (ASMS)–based HTS. By enriching true positives and filtering out non-binders upfront, HTS-Oracle streamlines the discovery pipeline and enables more focused, cost-effective screening. Two hit compounds disrupted the CD28–B7.1 interaction, with orthogonal validation provided by MST, ELISA, and molecular dynamics simulations. HTS-Oracle reduces screening burden and improves discovery efficiency, offering a powerful, scalable, and experimentally validated AI framework for accelerating hit identification across difficult-to-drug targets.

## 1. Introduction

High-throughput screening (HTS) remains a cornerstone of early-phase small molecule drug discovery, serving as the primary strategy for identifying initial chemical hits across therapeutic areas^1, 2, 3^. An analysis of 58 approved drugs showed that approximately 30% originated from HTS campaigns, and this approach remains the predominant method for hit identification in many leading pharmaceutical companies^4^. Over the years, HTS has undergone remarkable technological evolution, with innovative platforms such as temperature-related intensity change (TRIC)^5^, surface plasmon resonance (SPR)^6^, and mass spectrometry-based approaches^7^ enhancing the speed, resolution, and diversity of hit detection.

Despite technological advancements, conventional HTS campaigns often suffer from high costs, limited efficiency, and low success rates—typically yielding hit rates of just 1–2%^8, 9^. These inefficiencies are especially pronounced when screening against targets with limited tractability to small molecules, such as immune receptors and protein–protein interaction (PPI) interfaces. In addition to high attrition, HTS workflows often discard negative screening data, eliminating valuable information that could inform future predictions and compound prioritization^10, 11^. As a result, vast amounts of experimental data remain underutilized, and chemical space exploration remains largely unguided.

Artificial intelligence (AI) and machine learning (ML) offer a powerful opportunity to address these challenges by enabling large-scale virtual screening and compound prioritization^12^. Deep learning models have shown success in capturing nonlinear structure–activity relationships and accelerating hit discovery across vast chemical spaces^13, 14^. However, many ML frameworks are designed for retrospective performance benchmarking, lack integration with real-world HTS pipelines, and fail to incorporate negative data, limiting their prospective utility in challenging screening contexts. To overcome these limitations, we developed HTS-Oracle, a retrainable, multi-modal deep learning platform that integrates transformer-derived molecular embeddings with traditional cheminformatics descriptors. HTS-Oracle combines ChemBERTa^15^, a SMILES-based language model—with RDKit-derived molecular fingerprints and physicochemical features in an ensemble architecture optimized for prospective hit prediction. By uniting contextual and structural representations through multiple feature selection strategies, HTS-Oracle prioritizes high-confidence hits from large compound libraries. Unlike traditional Quantitative Structure– Activity Relationship (QSAR) models, HTS-Oracle is explicitly designed for prospective screening, retraining, and deployment across new targets—enabling rapid adaptation and reuse.

We benchmarked HTS-Oracle against CD28, a co-stimulatory immune receptor that plays a central role in T-cell activation and has long eluded small-molecule targeting due to its shallow and flexible extracellular interface^16, 17^. The interaction between CD28 and its ligand B7.1 (CD80) is a validated co-stimulatory axis relevant to autoimmune disease and transplant rejection ^18, 19, 20^. Although this interaction is successfully modulated by the fusion biologic CTLA-4-Ig (abatacept), no small molecule inhibitors of CD28 have reached clinical translation^21, 22^. We then applied HTS-Oracle to prospectively prioritize CD28-binding compounds from a chemically diverse small molecule library, enabling experimental evaluation of its predictive performance.

Experimental validation using TRIC identified 29 hits from 345 ML-prioritized compounds (8.4% hit rate), representing an eightfold improvement over traditional HTS methods including SPR, TRIC, and mass spectrometry–based approaches. Two compounds disrupted the CD28–B7.1 interaction with micromolar potency, supported by orthogonal validation via MST, competitive ELISA, and molecular dynamics simulations. These findings establish HTS-Oracle as a scalable, generalizable, and experimentally validated AI platform for efficient hit discovery—particularly against targets historically regarded as difficult to drug. Its prospective utility, retraining flexibility, and demonstrated enrichment over conventional HTS workflows offer a practical framework for accelerating early-stage drug discovery. A schematic overview of the HTS-Oracle framework—including model architecture, training and validation strategy, and application to CD28—is shown in Figure 1. Looking ahead, HTS-Oracle establishes a generalizable framework for integrating deep learning into prospective screening workflows, enabling efficient and scalable hit identification across structurally diverse and pharmacologically relevant targets.

**Figure 1.**
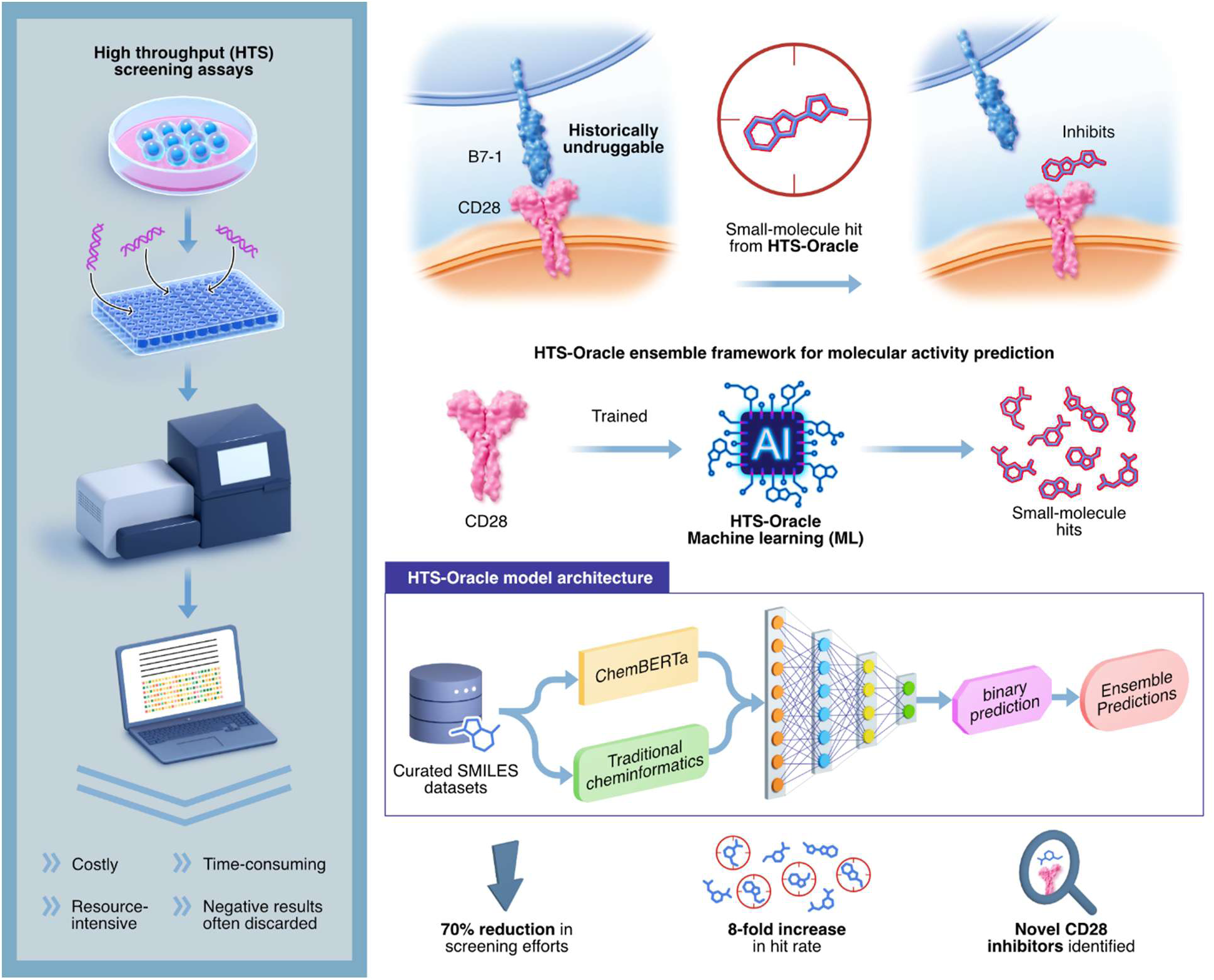
Schematic overview of the HTS-Oracle platform. Traditional HTS workflows are resource-intensive, time-consuming, and often discard negative data. HTS-Oracle addresses these limitations through a retrainable, multi-modal deep learning framework that integrates ChemBERTa embeddings with cheminformatics descriptors to predict compound activity. The model was trained on CD28 screening data and prospectively applied to prioritize candidate inhibitors from a chemically diverse library. Experimental validation via TRIC and orthogonal assays identified novel small molecule CD28 inhibitors, achieving an eightfold increase in hit rate and a 70% reduction in screening burden.

## 2. Results and Discussion

### 2.1. Training Data Engineering and Preprocessing

To develop a high-quality training dataset for ML–based prediction of CD28-binding small molecules, we previously optimized two orthogonal HTS platforms: a TRIC-based binding assay^23^ and a high-throughput mass spectrometry method for direct binder identification. Both assays targeted the extracellular domain of human CD28 (Asn19–Pro152), which was expressed with a His-tag for consistent orientation and physiological relevance.

The initial screening set consisted of over 7,000 curated small molecules from two chemically diverse libraries. Prior to model development, the compounds underwent rigorous preprocessing to ensure data integrity and modeling suitability. This included the removal of invalid SMILES strings, resolution of tautomeric and protonation inconsistencies, and elimination of structurally redundant molecules. These steps yielded a final non-redundant dataset of 6,003 unique compounds for model training. The TRIC and mass spectrometry assays collectively yielded 128 primary hits, which were subsequently refined through an additional round of curation. After removing invalid SMILES entries and duplicate structures, 120 unique compounds were retained as the positive class.

These 120 compounds were used as the “hit” category for supervised model training. While not all were confirmed in secondary validation assays, their inclusion provided a sufficiently large and chemically diverse positive dataset—a critical factor for enhancing model learning and generalizability across unseen chemical space. The remaining compounds in the dataset, which did not exhibit measurable CD28 binding in primary screens, were designated as negative examples.

Following model training and internal validation, we deployed the pipeline as a Streamlit-based web application, allowing accessible and user-friendly compound prioritization. A schematic overview of the dataset construction, curation pipeline, model architecture, and screening workflow is provided in Figure 2. All curated datasets and code used for training, validating, and evaluating the ML model are publicly available via our GitHub repository.

**Figure 2.**
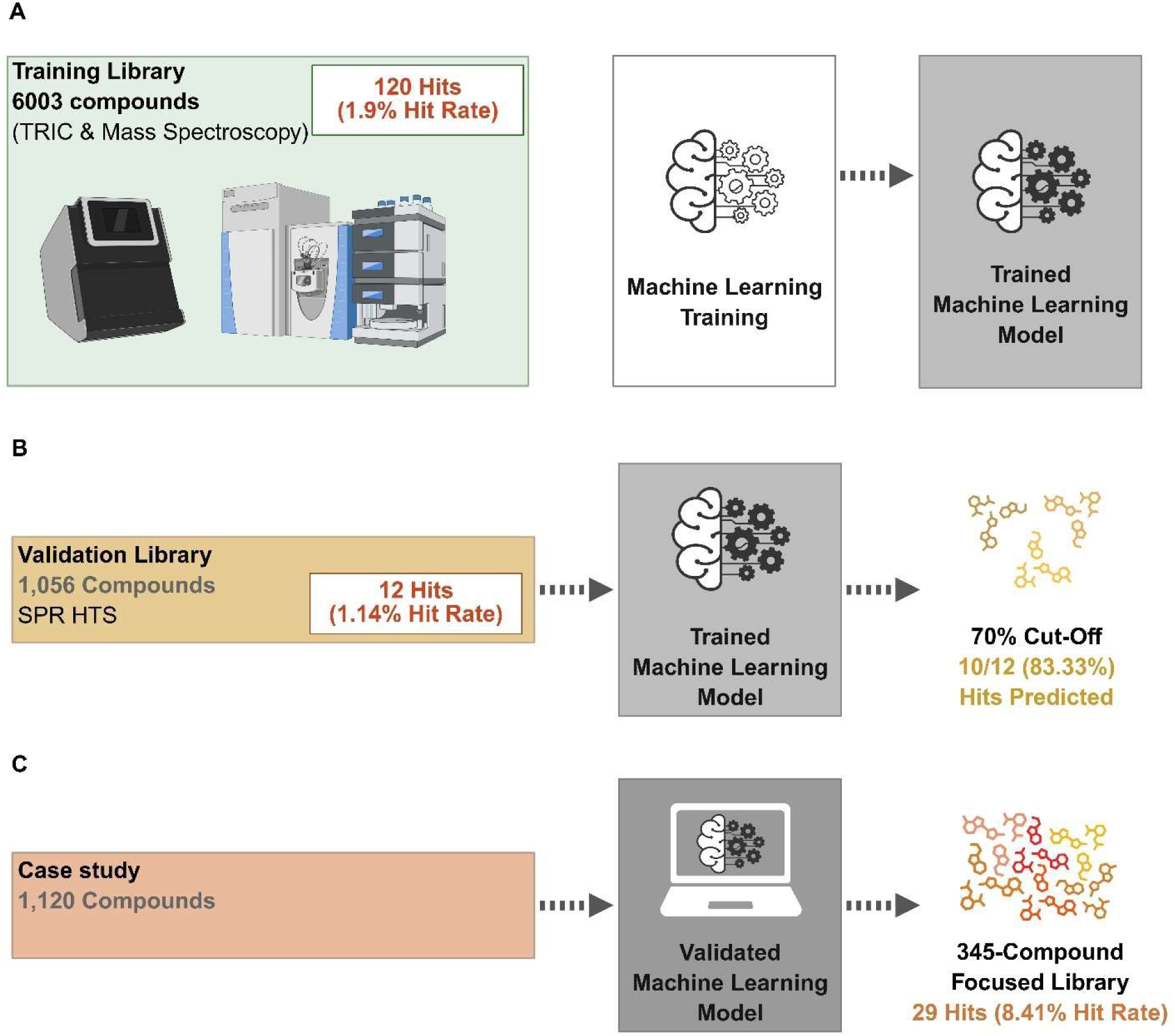
Schematic overview of the training, validation, and application of the ML model. **(A)** Training dataset consisting of 6,003 compounds with an observed hit rate of 1.9%. **(B)** Validation dataset containing 1,056 compounds with a hit rate of 1.14%. **(C)** Prospective application of the model on a novel compound library of 1,120 molecules, followed by experimental validation of the predicted hits, resulting in a significantly improved hit rate of 8.4%. (Created by Biorender)

To assess the generalizability of the trained model, we generated an independent *Validation Set* using a third HTS assay based on SPR^24^. This screen targeted the same extracellular domain of CD28, but utilized a biotinylated form of the protein for capture on a Sensor Chip CAP via Avitag™/Streptavidin coupling. This setup ensured stable ligand presentation across multiple regeneration cycles and enabled consistent signal quality. The SPR-based *Validation Set* consisted of 1,056 compounds from the Discovery Diversity Set (DDS, Enamine), a commercially available collection selected for scaffold diversity. The screen identified 12 hits (1.14% hit rate), which were not present in the original training set, ensuring that the validation was fully out-of-sample.

The trained ML model was applied to this dataset using a 70% prediction probability threshold and successfully identified 10 of the 12 confirmed hits, yielding a sensitivity of 83.3%. Based on this performance, the model was considered validated and suitable for deployment in prospective compound prioritization. We then applied the model to an *Evaluation Set* composed of 1,152 additional DDS compounds not present in either the training or validation sets. The model prioritized 345 molecules (30% of the set), enabling a ∼70% reduction in the number of compounds requiring experimental testing. Prospective screening of this focused library using TRIC identified 29 hits, corresponding to a hit rate of 8.41%—representing an eightfold enrichment over the historical baseline (1–2%) for this target.

Hit selection from this library via SPR was based on multi-parameter sensorgram analysis^24^. Key criteria included level of occupancy (LO) and binding kinetics, the latter of which excluded compounds showing non-specific binding (NSB) or non-dissociating behavior (NDB). NSB compounds were identified by elevated responses on the reference flow cell (higher than 5 response units), while NDB compounds were flagged based on minimal dissociation over time. Compounds with LO values above 200% were excluded, whereas those with LO between 50% and 100% and no NSB flag were prioritized as potential hits.

### 2.2. Model Architecture and Training

A hybrid deep learning model was constructed with a multi-modal ensemble architecture, designed to integrate chemical language modeling with traditional molecular descriptors for binary classification of chemical structures. At the core of this architecture (Figure 3) is a dual-branch neural network that processes two complementary types of molecular representations in parallel. The first branch leverages ChemBERTa^15^, a RoBERTa-based transformer model pre-trained specifically on chemical SMILES strings, to extract contextualized molecular embeddings. The ChemBERTa branch generates 768-dimensional embeddings from the classification [CLS] token, which are regularized using a dropout layer before being input to a multi-layer perceptron (MLP) composed of two linear layers (768→256→128) with batch normalization, ReLU activation, and 30% dropout between layers.

**Figure 3:**
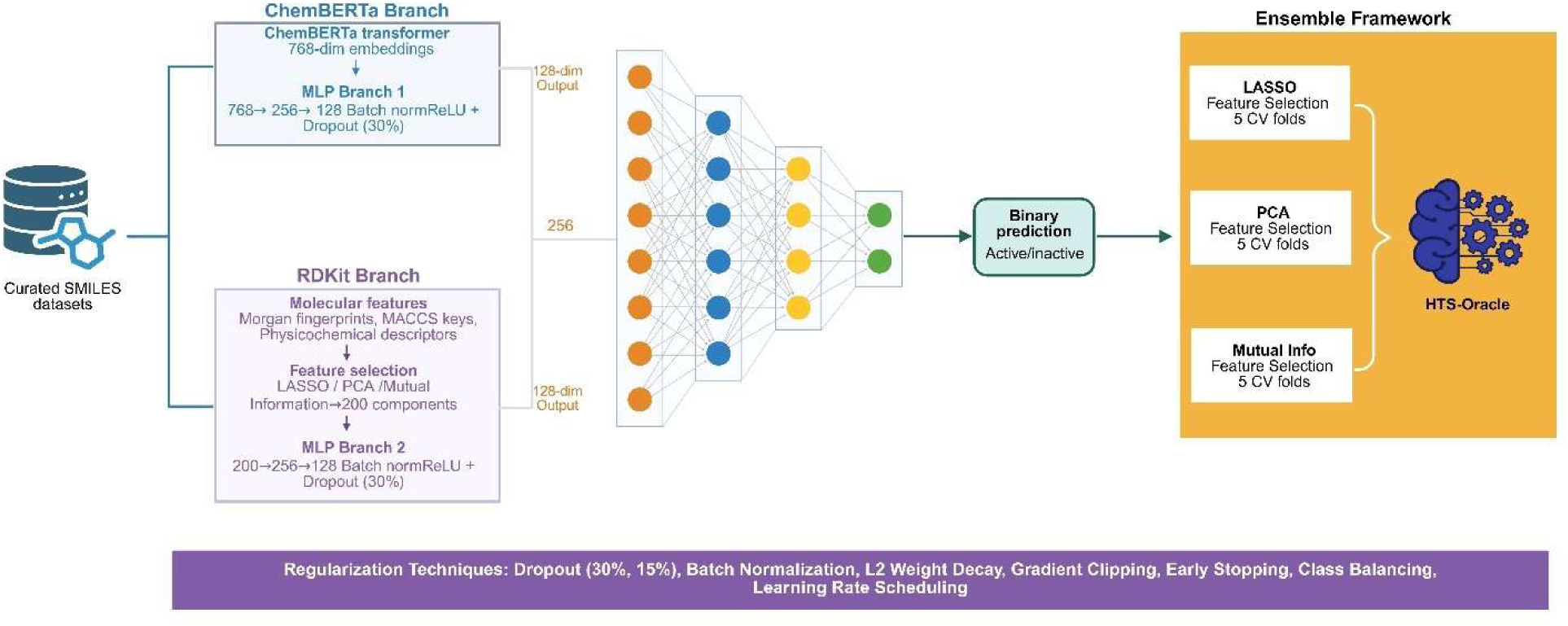
Architecture of the Hybrid Ensemble Model for Molecular Property Prediction (Created by Biorender). The model features two parallel processing branches that merge via late fusion: (A) ChemBERTa Branch (Top): Tokenized SMILES strings are processed through ChemBERTa’s embedding layer and 12 transformer layers. The resulting CLS token embedding (768-dim) then passes through two hidden layers (256 → 128-dim), each incorporating dropout (p=0.3) and batch normalization. (B) RDKit Branch (Bottom): Engineered molecular features (2,230-dim, including Morgan fingerprints, MACCS keys, and physicochemical descriptors) are fed into this branch. These features undergo optional dimensionality reduction via LASSO, PCA, or mutual information before passing through two hidden layers (256 → 128-dim) with dropout and batch normalization. The outputs from both branches are concatenated (forming a 256-dim combined feature vector) and passed to a three-layer classifier (128 → 64 → 1-dim) with a sigmoid activation function for final prediction. An ensemble model is then formed by aggregating predictions from the different feature selection methods (LASSO, PCA, mutual information) using mean voting.

The second branch handles cheminformatics features derived from RDKit^25^ which included 2048-bit Morgan fingerprints, 167-bit MACCS keys, and 15 physicochemical descriptors (e.g., molecular weight, Log*P*, rotatable bonds, hydrogen bond donors and acceptors, topological polar surface area, ring systems, and drug-likeness metrics). These features undergo dimensionality reduction using one of three methods—LASSO-based feature selection, principal component analysis (PCA), or mutual information filtering which typically reduces the feature space to 200 components. This reduced representation is passed through a parallel MLP with an architecture identical to that of the ChemBERTa branch, allowing harmonized processing of diverse input modalities.

The 128-dimensional outputs from both branches were concatenated into a unified 256-dimensional representation and passed through a final classification head comprising three fully connected layers (256→128→64→1). This classifier used batch normalization and ReLU activations between layers and applied graduated dropout rates (30% and 15%) to encourage generalization and mitigate overfitting. The model employed an ensemble framework by training distinct models for each feature selection method (LASSO, PCA, mutual information) across five stratified folds, yielding a total of 15 sub-models. During inference, predictions from all sub-models were averaged to produce the final ensemble output, enhancing robustness and minimizing overfitting.

To prevent overfitting and minimize bias, the model incorporated multiple regularization techniques, including dropout at multiple stages, batch normalization to reduce internal covariate shift^26^, L2 weight decay *via* the AdamW optimizer, and gradient clipping to stabilize training^27^. Early stopping was employed with a five-epoch patience window based on validation AUC to avoid overtraining. To address the class imbalance inherent in HTS datasets, where hits are sparse, positive class weights were dynamically balanced^28^ for the binary cross-entropy loss based on the class distribution in each training fold. This approach prevented the model from favoring the majority class.

The training pipeline also included several mechanisms to ensure data integrity and model robustness. SMILES strings were validated and parsed with error-handling fallbacks, while molecular features were standardized using StandardScaler to normalize scales across descriptors. NaN and infinite values were automatically replaced with zeros, and safe fallback strategies were used during metric computation to handle edge cases such as single-class predictions. Learning rate scheduling was handled *via* a ReduceLROnPlateau strategy^29^, which adapted the learning rate based on validation performance to avoid premature convergence or overshooting. Finally, the integrity of the validation step was maintained through stratified train–validation splits^30^, ensuring balanced class distributions without leakage. Each fold had an independent validation set to provide unbiased performance estimates.

Training of the model demonstrated an AUC-ROC of 0.9914 and an average precision of 0.8565 (Figure 4). The three feature selection methods showed varying individual performance levels. PCA achieved the highest individual AUC-ROC of 0.9922 with an average precision of 0.8608, closely matching the ensemble performance. The mutual information method demonstrated strong performance with an AUC-ROC of 0.9792 and average precision of 0.7140. LASSO showed an AUC-ROC of 0.8846 and average precision of 0.6069. The ensemble’s superior performance over individual methods is evident in the ROC curve comparisons, where the ensemble curve (AUC = 0.9914) outperformed all individual methods. This improvement demonstrates the benefit of combining complementary feature selection approaches, as each method captures different aspects of the feature space.

**Figure 4.**
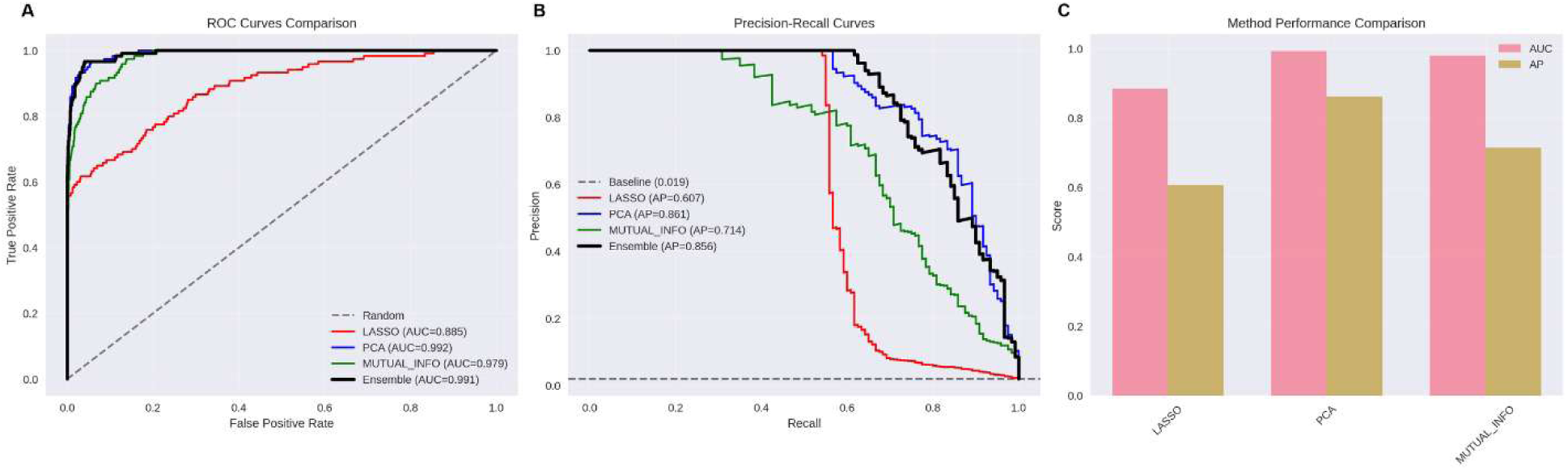
ROC and Precision-Recall Curves Comparison. (A) ROC curves comparing ensemble performance (black line, AUC = 0.991) against individual methods: LASSO (red, AUC = 0.885), PCA (blue, AUC = 0.992), and MUTUAL_INFO (green, AUC = 0.979). Diagonal dashed line represents random performance. (B) Precision-recall curves showing ensemble (black, AP = 0.856) versus individual methods with corresponding average precision scores. (C) Performance comparison bar chart displaying AUC-ROC (pink bars) and average precision (brown bars) for each method.

Further evaluation of the trained model’s performance showed that there was a strong correlation between predictions generated using LASSO, PCA, and Mutual Information for feature selection, with all pairwise correlations ≥0.75 (Figure 5A). This high level of agreement across diverse dimensionality reduction techniques was indicative of consistent signal capture which reinforces the reliability of the ensemble approach. Further supporting this, the “Cross-Validation Performance” panel (Figure 5E) showed that all methods achieved similarly high validation AUCs (∼0.80–0.85) with narrow interquartile ranges, demonstrating strong generalizability across folds. The “Enrichment in Top Predictions” bar chart (Figure 5C) highlights the model’s ability to prioritize true binders effectively, achieving 53-fold enrichment in the top 1% of predictions. Although enrichment decreased for larger ranked subsets, it consistently remained well above random expectation, confirming the model’s value in guiding experimental screening.

**Figure 5.**
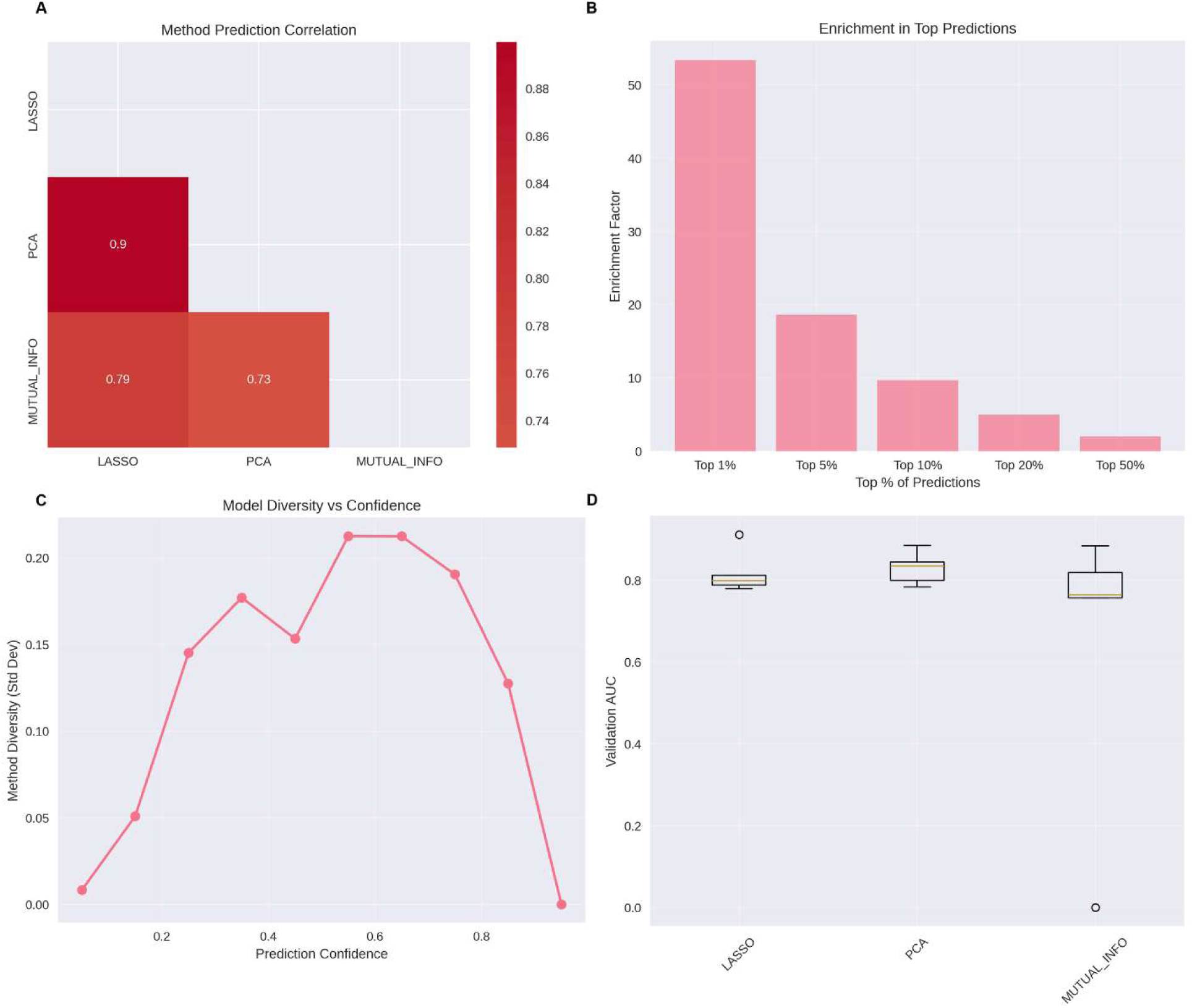
Performance Evaluation of the Hybrid Ensemble Model. (A) Method Prediction Correlation: Heatmap showing pairwise correlation coefficients between predictions generated using different feature selection methods (LASSO, PCA, and Mutual Information). (B) Enrichment in Top Predictions: Bar chart illustrating the enrichment factor of true positive hits within the top-ranked prediction subsets (Top 1% to 50%). (C) Model Diversity vs. Confidence: Plot of the standard deviation in predicted probabilities across feature selection methods as a function of prediction confidence. (D) Cross-Validation Performance: Box plots showing the distribution of validation AUC scores for models trained with each feature selection method.

Prediction confidence was found to align closely with both model consensus and predictive accuracy. As shown in the “Model Diversity vs. Confidence” plot (Figure 5D), diversity among model predictions is lowest at the confidence extremes which is an indicator of strong consensus. Meanwhile, the model predictions exhibited peaks between 0.5 and 0.7, near the decision boundary where predictions are most uncertain. Based on these observed patterns, along with consistent model agreement, high validation performance, and strong enrichment in top predictions (Figure 5), a confidence threshold of 0.7 was selected for inference on new datasets. This threshold strikes a balance between predictive reliability and practical prioritization, ensuring that high-confidence predictions reflect consensus-driven and experimentally actionable candidates. Together, the panels in Figure 5 demonstrate that the ensemble model is robust, well-calibrated, and consistent. Its ability to integrate diverse feature sets while maintaining high predictive performance highlights its value as an effective tool for prioritizing candidate binders and streamlining hit discovery.

Next, 3D PCA visualization (Figure 6) was carried out to understand how the trained model behaves within the multidimensional chemical space of the molecular dataset. The PCA visualization showed that the training library was well distributed in a roughly spherical cluster within the first three principal components, which is characteristic of well-curated molecular datasets. Moreover, the true positive compounds (initial hits) were well distributed throughout this space (Figure 6A) rather than clustering in distinct regions, indicating that bioactivity is not confined to specific structural scaffolds or chemical families, further validating the chosen training dataset.

**Figure 6.**
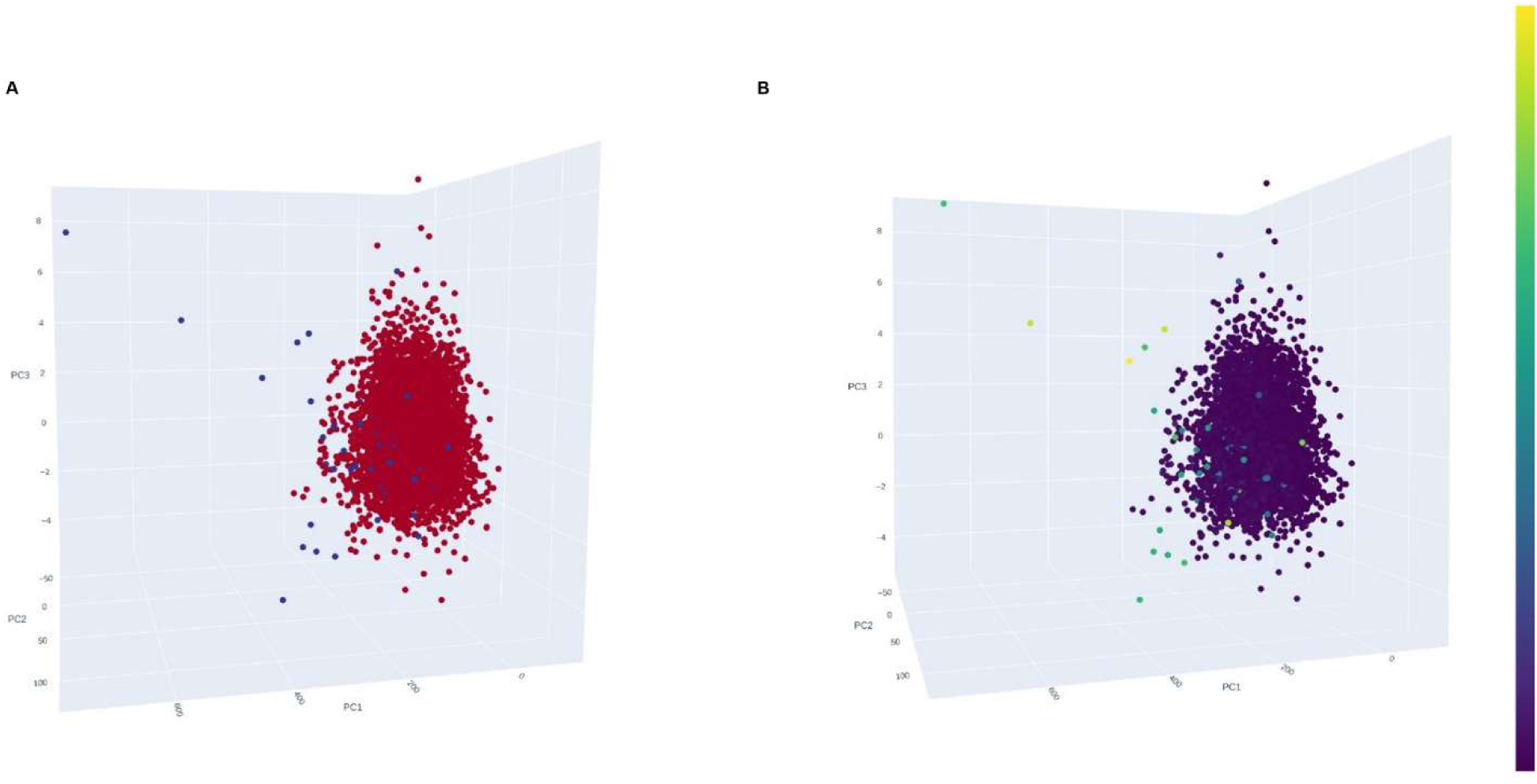
3D Principal Component Analysis of Molecular Chemical Space. The visualization shows the distribution of molecules projected onto the first three principal components (PC1, PC2, PC3) of the molecular descriptor space. (Left panel) True labels are displayed with active compounds shown in red and inactive compounds in blue. (Right panel) Predicted labels are shown with color intensity representing model prediction scores, where dark purple indicates predicted inactives (score ∼0) and bright yellow indicates predicted actives (score ∼1). Each point represents a single molecule positioned according to its chemical similarity relationships. The 3D coordinate system reveals the natural clustering structure of the chemical space, with molecules distributed in a roughly spherical pattern typical of drug-like compound libraries. Active compounds (true positives) are dispersed throughout the chemical space rather than concentrated in specific regions, indicating diverse structure-activity relationships.

The predicted labels visualization demonstrated that the trained model maintained a consistent performance characteristic across the entire chemical space. The prediction confidence scores, represented by the color gradient from purple (predicted negatives) to yellow (predicted positives) shows that the predicted hits are distributed throughout the space rather than concentrated in specific regions in a similar fashion to the library. This pattern indicates that the model has learned generalizable structure-activity relationships that are not biased toward particular chemical scaffolds or structural classes. These observations support the ability of model to identify active compounds from novel chemical scaffold and its ability to capture fundamental structure-activity principles that transcend simple chemical similarity. This characteristic is particularly valuable for discovering new chemical starting points and expanding the chemical diversity of compound libraries.

### 2.3. HTS-Oracle

A Streamlit application, named HTS-Oracle, was developed to provide an interactive and scalable platform for predicting molecular activity across compound libraries using the pre-trained ensemble model. At its core, the application integrates multiple feature selection methods (LASSO, PCA, and mutual information) to generate calibrated activity scores for each compound, offering users not only binary classifications but also nuanced confidence levels.

In scenarios where no known actives are available to train a predictive model, the HTS-Oracle application includes a robust backup prediction system that filters libraries based on classical drug-likeness rules. This system applies Lipinski’s Rule of Five and Quantitative Estimate of Drug-likeness (QED) to rank the compounds based on their predicted absorption, distribution, and overall bioavailability profiles.

Beyond predictions, the application offers a suite of interactive visualizations powered by Plotly and Matplotlib, enabling users to explore the distribution of prediction scores, compare feature selection methods, and correlate predicted activity with key molecular properties. With built-in error handling for invalid SMILES, support for model interpretability, and configurable thresholding, the app serves as a practical tool for virtual screening, hit prioritization, and rational library design in early-stage drug discovery.

### 2.4. ML Model Validation

To assess the predictive performance of our trained model on an independent dataset, we employed a dataset based on the results of a HTS campaign utilizing surface plasmon resonance (SPR) technology recently conducted by our laboratory.^24^ This validation approach ensures that model performance evaluation is conducted on truly unseen data, thereby providing an unbiased assessment of generalizability.

The validation dataset comprised 1,056 structurally diverse compounds selected from the Discovery Diversity Set (DDS) library supplied by Enamine. The SPR-based HTS assay identified 12 compounds as primary hits that exhibited binding affinity to CD28, corresponding to a hit rate of 1.14%.

The same 1,056-compound library was subjected to a screening using our HTS-Oracle pre-trained model deployed through a Streamlit web application interface. At an 80% prediction confidence cutoff, the HTS-Oracle model successfully retained 8 out of the 12 experimentally-confirmed hits (66.7% recall) within the top 20% of ranked compounds. Meanwhile, when the confidence threshold was relaxed to 70%, the performance of HTS-Oracle improved significantly, capturing 10 out of the 12 validated hits (83.3% recall) within the top 30% of compounds. The 83.3% recall rate at this threshold suggests that the model effectively captures the majority of experimentally-validated CD28 binders, making it a valuable tool for prospective virtual screening campaigns.

Based on these comparative results and earlier observations, the 70% confidence threshold was selected for the planned evaluation screening assay. This decision balances the competing objectives of maximizing hit recovery (recall) while maintaining reasonable prediction confidence. Moreover, the validation results instill confidence in our chosen approach and supports its utility as a complementary tool to traditional high-throughput screening methodologies.

### 2.5. Translational Evaluation and Deployment for CD28 Binder Identification

To evaluate the predictive performance of the ML model trained on CD28-binding compounds, we applied it to an independent small molecule library with the goal of enriching for true CD28 binders while reducing the number of compounds requiring experimental screening. The extracellular domain of the His-tagged human CD28 protein (Asn19–Pro152) was used as the assay target, consistent with previous screening campaigns.

To identify potential CD28 binders, we began by selecting a diverse collection of 1,120 small molecules from Enamine’s Discovery Diversity Set which served as the input screening library for our trained ML model. We then applied our model to this library with compounds flagged as potential binders if their prediction probability exceeded a 70% threshold. This rigorous filtering process allowed us to narrow the initial large set to yield a focused subset of 345 compounds that were predicted to have a high likelihood of binding to the CD28 protein. This targeted approach significantly reduced the number of compounds requiring further experimental validation, streamlining the discovery process for novel CD28 binders (Figure 2).

#### 2.6.1. TRIC-based affinity screening

Compounds were screened in 384-well plates at a concentration of 200 µM using a PBS-based assay buffer. Hit selection was based on ΔF_norm_ values derived from TRIC measurements and compounds exceeding three times the standard deviation of the negative control were considered to be hits. Compounds exhibiting autofluorescence, quenching, or aggregation artifacts were excluded.

Based on these selection criteria, 29 primary hits were identified from the computationally focused library of 345 compounds, yielding a remarkably high hit rate of 8.41% (Figure 7). When benchmarked against the original unfiltered library of 1,120 compounds, our trained model achieved a dramatic 70% reduction in the experimental screening effort while simultaneously delivering an 8-fold enhancement in hit rate compared to previous CD28-targeted HTS campaigns (Figure 2). The 8.41% hit rate achieved through this focused screening represents a marked departure from the typical 1%^8, 31^ hit rates commonly observed in HTS campaigns.

**Figure 7.**
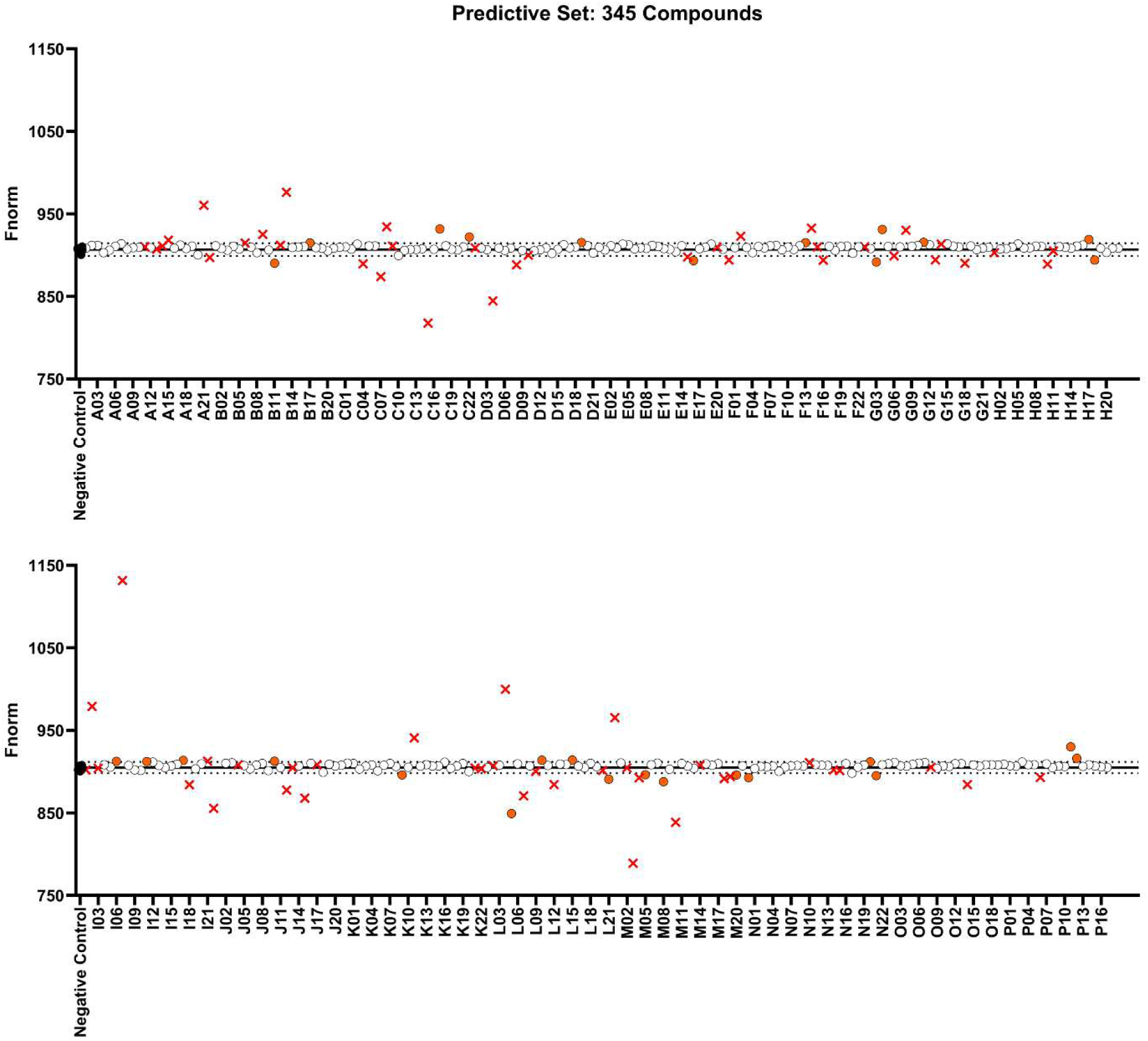
ML-Guided HTS Using TRIC Reveals CD28 Binders from the Discovery Diversity Set. Each data point represents the normalized fluorescence value (F_norm_) for an individual compound screened at 200 µM. Two sets of 16 negative control samples (black dots) were included on each half of the 384-well plate, corresponding to the two acquisition batches (top and bottom halves of the 384-well plate). The mean of the negative controls (solid black line) and their standard deviation were used to define the threshold for identifying potential CD28 binders. This threshold was defined as a response amplitude (ΔF_norm_) exceeding three times the standard deviation of the negative control (black dotted lines). Compounds exhibiting scan anomalies, autofluorescence, quenching of the RED-tris-NTA dye, or aggregation were excluded and are marked with red crosses. The remaining 29 compounds (orange dots) above or below the threshold lines were identified as potential CD28 binders from the 345-compound ML–prioritized library. Data shown are from a single HTS experiment performed using the Dianthus instrument (NanoTemper Technologies).

The improved performance of this focused approach highlights the effectiveness of our trained model and chosen approach in reducing the number of compounds that need to be screened, thereby conserving resources, and in significantly increasing the likelihood of identifying viable lead compounds. This gain in efficiency is especially notable given the historical challenges associated with targeting CD28.

#### 2.6.2. CD28 Binding Affinity Screening Using Spectral Shift Assay

Among the 29 hits identified in the TRIC-based screening, 14 compounds were commercially available and subsequently selected for binding affinity measurements using spectral shift detection. Of these, five compounds showed dose-dependent binding affinity toward CD28. Notably, three compounds exhibited quantifiable binding interactions with the CD28 protein (Figure 8). Compound Z2770993038 demonstrated the highest affinity, with a dissociation constant (K_d_) of 43.35 ± 16.44 μM, followed closely by Z4390463985 (*K*_d_ = 45.76 ± 12.06 μM) and Z3889890124 (K_d_ = 49.53 ± 15.3 μM). All three compounds produced characteristic dose-dependent binding curves with saturation behavior, confirming specific and saturable interactions with CD28 (Figure 8). Conversely, Z1558583902 and Z1896624998 showed limited binding responses within the tested concentration range, suggesting potential binding interactions that may occur at higher concentrations.

**Figure 8.**
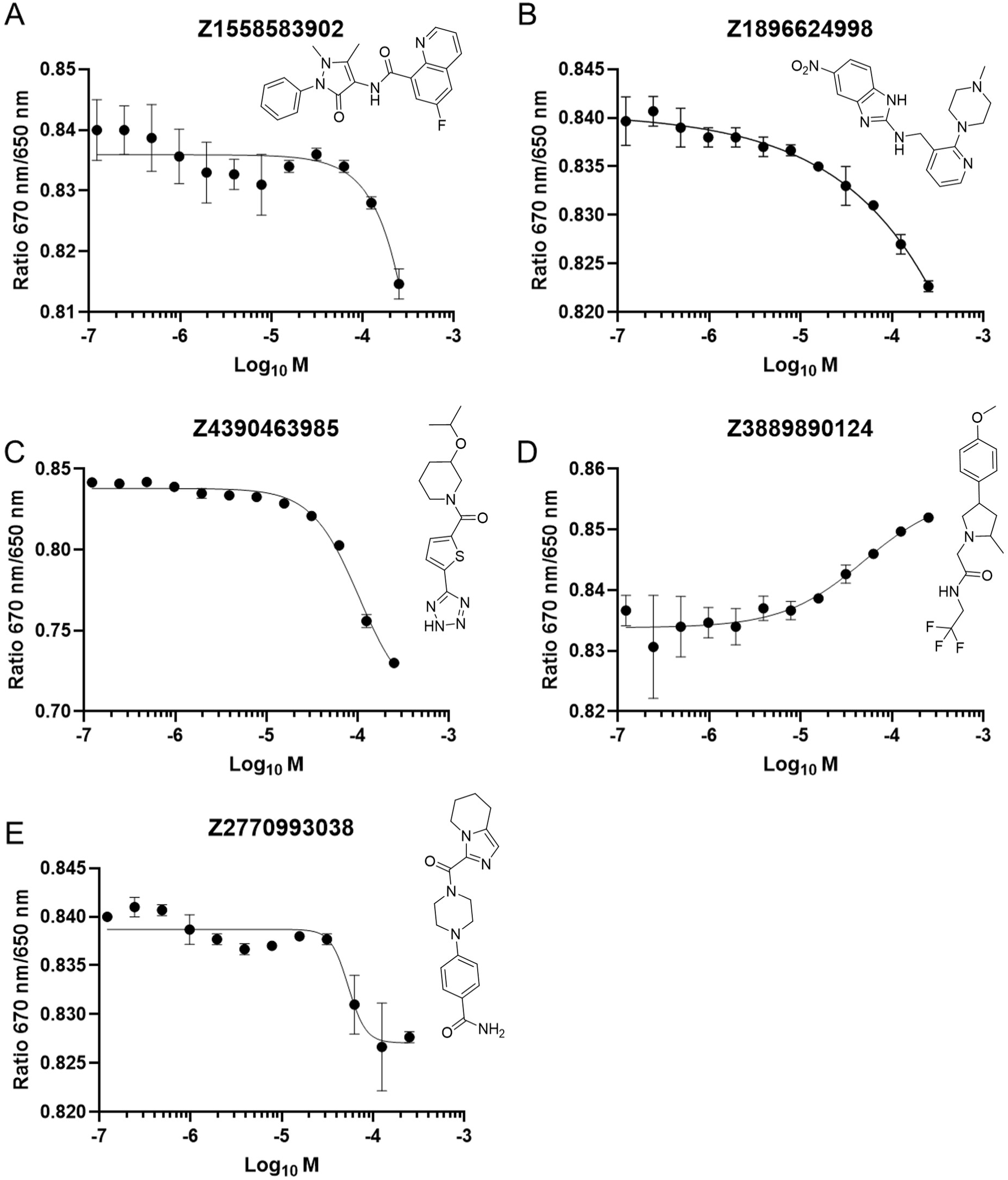
Binding affinity measurements of test compounds to CD28 protein using spectral shift assay. (A-E) Dose-response curves showing the binding of five test compounds to recombinant human CD28 protein (40 nM) measured by spectral shift detection using the Monolith X platform. The y-axis represents the ratio of fluorescence at 670 nm to 650 nm, and the x-axis shows the log₁₀ molar concentration of each compound. Data points represent mean ± SEM from three independent experiments. Binding curves were fitted using a one-site binding model in GraphPad Prism. The 2D structures of the tested compounds are overlayed on the top of each curve.

#### 2.6.3. Competitive ELISA assay

Targeting the CD28–B7.1 interaction represents a compelling therapeutic approach to modulate T cell co-stimulation, particularly in the context of autoimmune diseases, transplant rejection, and cancer ^32, 33, 34^. CD28 delivers essential co-stimulatory signals that enhance T cell proliferation and drive the secretion of pro-inflammatory cytokines such as IL-2, IFN-γ, and TNF-α^35, 36, 37^. To investigate the ability of small molecules to block CD28–B7.1 binding, we performed a competitive ELISA using immobilized CD28 and biotinylated B7.1 ^24^. Two compounds, Z2770993038 and Z4390463985, were evaluated across a concentration range. Both compounds demonstrated dose-dependent inhibition of CD28-B7.1 binding, with IC₅₀ values of 32.87 ± 5.74 µM and 56.29 ± 11.24 µM, respectively (Figure 9). These functional results correlate with their binding affinity obtained from the MST assay, confirming them as CD28 binders. Overall, these findings confirm that both compounds directly target CD28 and can disrupt its interaction with B7.1, supporting their potential as early-stage small-molecule inhibitors for further lead optimization and therapeutic development.

**Figure 9.**
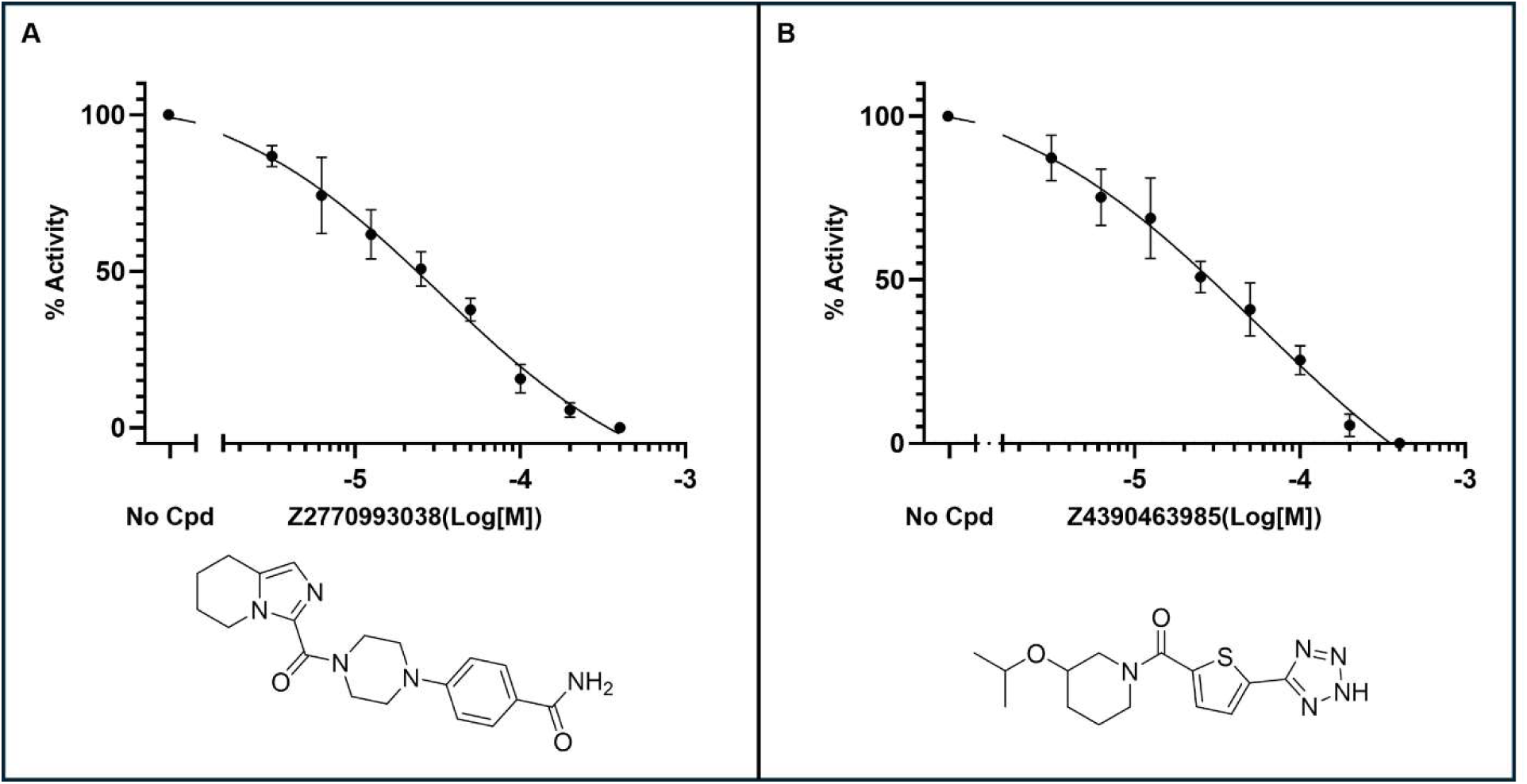
Dose-dependent inhibition of CD28–B7.1 interaction by compounds. The ability of compounds, Z2770993038 and Z4390463985 to inhibit CD28–B7.1 binding was assessed using a competitive ELISA assay. Z2770993038 and Z4390463985 exhibited a dose-dependent effect, with an IC₅₀ of 32.87 ± 5.74 µM and 56.29 ± 11.24 µM, respectively. Relative activity was calculated by normalizing luminescence values, setting the no-compound signal as 100% and the maximal compound-treated signal as 0%; all intermediate values were normalized accordingly. IC₅₀ was determined using GraphPad Prism with a four-parameter variable slope nonlinear regression model. Data represent mean ± standard deviation from 3 independent repeats.

### 2.6. In-silico Investigation of the Top Hits

The binding modes and stability of CD28-ligand complexes were analyzed for the two top hit compounds Z2770993038 and Z4390463985 in an effort to understand their mechanism of action. The compounds were docked into the secondary binding pocket of CD28 which was previously identified as possessing favorable features for small molecule targeting^24^. Both compounds were predicted to establish one hydrogen-bond with amino acid residues in the binding site as well as engaging in several hydrophobic interactions. The hydrogen-bond established by the primary amide of Z2770993038 was with the backbone carbonyl of Ala44 while the carbonyl group of Z4390463985 established its hydrogen-bond with the amine group (side chain ammonium) of Lys95 via the (Figure 10A-D). Interestingly, the presence of an S–H···π interaction between the thiophene moiety of Z4390463985 and the backbone hydrogen of His38 is proposed to facilitate the alignment of the compound with Lys95, thereby stabilizing the predicted hydrogen bond.

**Figure 10.**
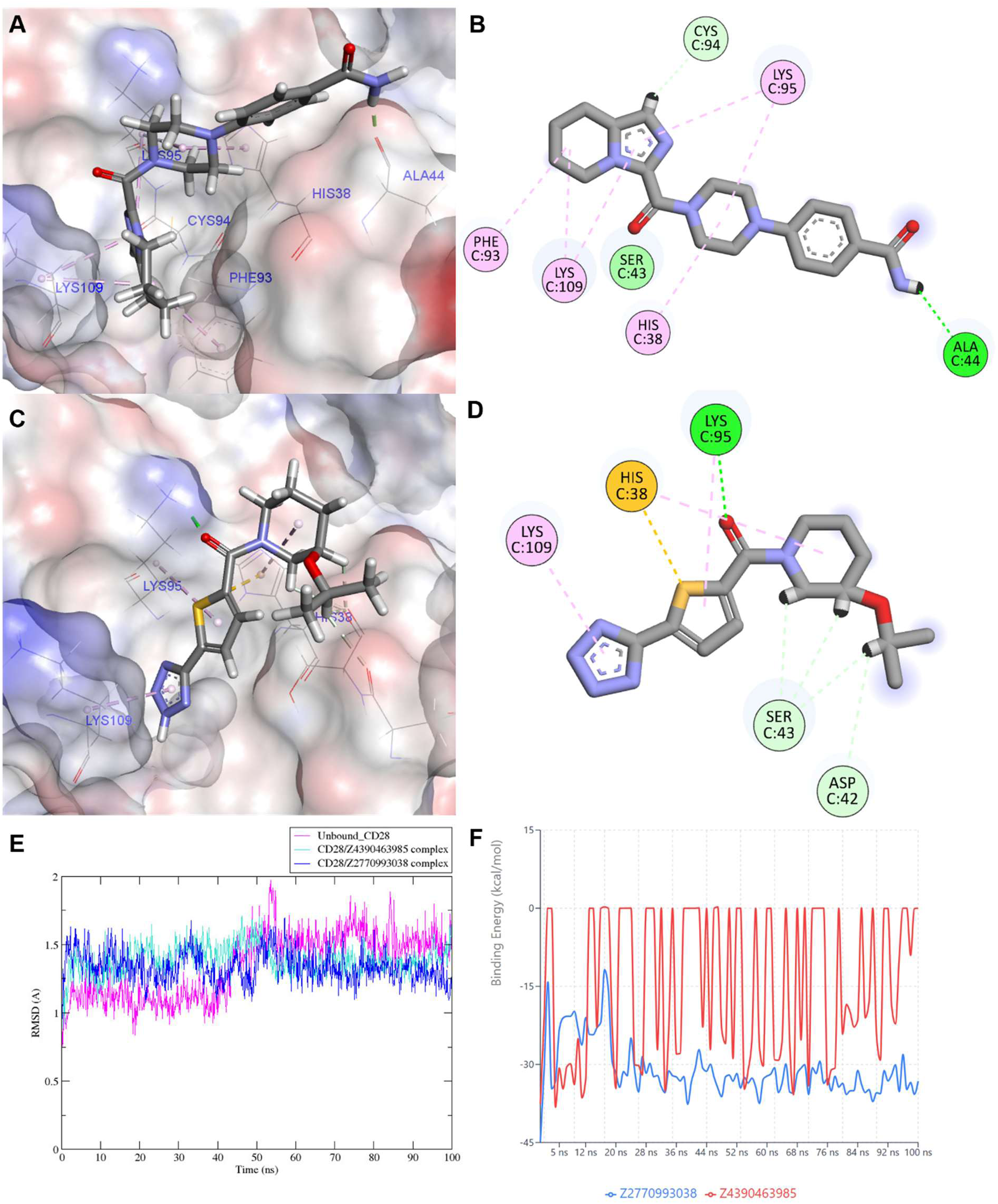
In-silico prediction of the binding modes of Z2770993038 and Z4390463985 with CD28. (A, B) 3D Docked pose and 2D interaction diagram of Z2770993038 within the CD28 binding pocket. (C, D) 3D Docked pose and 2D interaction diagram of Z4390463985 within the same site. Key interacting residues are labeled. (E) RMSD plots over 100 ns molecular dynamics simulations comparing unbound CD28, CD28/Z2770993038, and CD28/Z4390463985 complexes. (F) Binding energy fluctuations over time derived from MMGBSA calculations for both ligand–receptor complexes. Z4390463985 shows stronger and more stable binding.

Three 100 ns molecular dynamics (MD) simulations were conducted to compare the structural and energetic stability of Z2770993038 and Z4390463985 with CD28 with that of the unbound CD28 protein (Figure 10E–F). Analysis of root-mean-square deviation (RMSD) trajectories (Figure 10E) revealed that both ligand-bound systems conferred greater structural stability to the CD28 protein when compared to the unbound form. Notably, the Z2770993038/CD28 complex exhibited a slightly lower average RMSD than the Z4390463985/CD28 complex, consistent with the higher experimental binding affinity observed for Z2770993038. Similarly, MMGBSA-based binding free energy calculations (Figure 10F) demonstrated that Z2770993038 established a more stable complex with CD28, with an average binding free energy of –31.14 kcal/mol, compared to –15.56 kcal/mol for Z4390463985. The observed difference in binding free energy of the two complexes can be explained by examining Figure 10F which shows that the free energy of Z4390463985/CD28 complex fluctuates significantly throughout the MD simulation while that of the Z2770993038/CD28 complex remains largely stable. The correlation between the experimental binding affinity and predicted energy profiles reinforces the potential of the identified hits and provides a mechanistic explanation for their proposed binding modes.

## 3. Methods

### 3.1. Model Development

A hybrid deep learning framework was developed to identify active compounds from HTS libraries by combining chemical structure-based representations with ensemble classification. Compounds were obtained from library.csv (containing SMILES strings and compound IDs) and positives.csv (containing confirmed actives). Each compound was assigned a binary activity label, where y=1, y=1 if its ID appeared in the positives set and y=0, otherwise.

Each SMILES string was featurized using three complementary molecular representations: Morgan circular fingerprints^38^ were computed with radius 2 and 2,048 bits, encoding neighborhood substructures. In parallel, 167-bit MACCS keys^39^ were generated to capture common substructural motifs. Additionally, a set of 15 physicochemical descriptors was computed using RDKit^25^, including molecular weight, Log*P*, topological polar surface area, hydrogen bond donors and acceptors, rotatable bonds, ring count, and QED. These vectors were concatenated to form a 2,230-dimensional feature vector per compound.

To reduce dimensionality, we applied three feature selection methods independently: LASSO regression^40^, principal component analysis (PCA)^41^, and mutual information ranking^42^. LASSO performs feature selection by minimizing the L1-penalized loss:

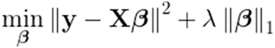

For PCA, features were projected onto principal components, and the top 200 were retained. Mutual information selection ranked features by their dependency with the activity labels. Subsequently, a dual-branch neural network was trained using five-fold stratified cross-validation. One branch used ChemBERTa^15^, a transformer-based model pretrained on SMILES, to encode each molecule’s SMILES into a dense vector representation. The second branch processed the selected RDKit features through a multilayer perceptron. Outputs from both branches were concatenated and passed through fully connected layers to produce a final prediction score. Training used the AdamW optimizer and binary cross-entropy loss with class weights to address data imbalance:

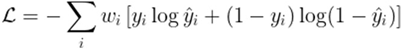

where *ŷ_i_* is the predicted probability and *w_i_* is the class weight.

Model predictions were averaged across folds and feature selection strategies. For each compound *i*, the final ensemble prediction was computed as the mean of its valid model outputs:

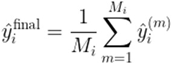

Model performance was assessed using the area under the receiver operating characteristic curve (ROC AUC), average precision (AP), precision, recall, and F1-score among other features. The script employed to evaluate the trained ML model “evaluation.py” is available in the GitHub repository. Final prediction scores were exported for downstream prioritization.

### 3.2. Streamlit Application

A Streamlit application was developed to implement the trained ensemble model for predicting molecular activity in new datasets, offering a user-friendly interface for researchers. The application allows users to upload molecular libraries in CSV format and receive predictions in a streamlined workflow. To ensure compatibility with diverse data sources encountered in chemical and pharmaceutical research, the application automatically identifies SMILES columns through pattern matching against standard naming conventions. For each molecular input, the application constructs a comprehensive descriptor-based representation. This feature generation step includes Morgan fingerprints, MACCS keys, and physicochemical descriptors. Morgan fingerprints are calculated from the SMILES representation using the equation:

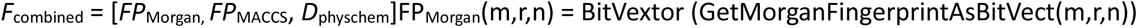

where *m* is the molecular structure, *r* = 2 the radius, and *n* = 2048 n=2048 is the fingerprint bit length. The final feature vector contains molecular properties such as molecular weight, Log*P*, rotatable bonds, hydrogen bond acceptors/donors, topological polar surface area, aromatic rings, heteroatoms, sp³ carbon fraction, and QED score.

The application uses an ensemble-based classification framework built on multiple ML models trained with different feature selection strategies, namely LASSO, PCA, and mutual information. For each method *k*, predictions are averaged across models using:

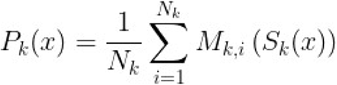

where *M_k,i_* is the *i^th^* model for method *k*, and *S_k_(x)* is the feature-selected transformation of the input vector *x*. The final ensemble prediction is obtained by averaging over all methods:

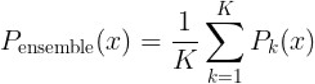

with K=3 in. Predictions are then converted into binary activity labels using a classification threshold θ (default = 0.5), defined as:

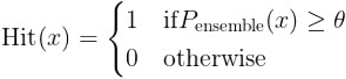

In addition to classification, the model also outputs prediction confidence, calculated as 2⋅∣P_ensemble_ (x) −0.5 ∣, and ensures that all scores are clipped to the range [0,1] to maintain probabilistic interpretability. To ensure reliability, SMILES validation is performed using RDKit parsing before feature extraction^43^. Descriptors are also sanitized by replacing NaN or infinite values with zeroes. The application supports efficient batch processing with real-time progress updates and robust error tracking.

Finally, prediction scores are visualized using interactive, color-coded tables, where color intensity scales linearly with predicted activity. This visualization aids users in rapidly identifying top candidates for downstream experimental validation.

### 3.3. Screened Libraries

To develop and validate a ML workflow for the identification of small-molecule binders to the human CD28 protein, a series of chemically diverse compound libraries were employed across training, validation, and prediction phases. The training set comprised of compounds which have been tested for CD28 small molecule binders identification using TRIC and Mass screening HTS technologies. The compound library were organized into two separate CSV files: “library.csv”, which contains the complete set of screened compounds, and “positives.csv”, which lists the subset of hits. Hits were defined as compounds that demonstrated activity in the primary single-dose HTS assays. Notably, these hits were not required to be confirmed through follow-up dose-response validation. This approach was deliberately chosen to maximize the number of positive examples available for training the ML model, acknowledging that some false positives may be present but prioritizing sufficient data coverage.

For model validation, a subset of 1,056 compounds was selected from the *Discovery Diversity Set* (Cat# DDS-10-10-Y-10, Enamine). These compounds were similarly formatted in 384-well plates, dissolved in DMSO at 10 mM, and stored at −80 °C. To assess the predictive performance of the trained model, a distinct subset of 1,152 compounds from the same *Discovery Diversity Set*, not overlapping with the validation set, was used as the final Evaluation Set. These compounds were handled under identical conditions.

Following HTS of the Evaluation Set, selected hit compounds were re-ordered as lyophilized powders from Enamine. These were reconstituted in DMSO at a 50 mM stock concentration and stored at −30 °C for downstream experimental validation.

### 3.4. High-Throughput Screening and Hit Selection for the Training Set Using TRIC Technology

HTS of the *Training Set* was performed using the Dianthus NT.23 Pico system (NanoTemper Technologies) as previously described in detail^23^. Briefly, human CD28 protein (His-tagged, Cat# CD8-H52H3, Acro Biosystems, Newark, DE, USA) was labeled with RED-tris-NTA dye using the Monolith His-Tag Labeling Kit (Cat# MO-L018, NanoTemper Technologies, München, Germany), following the manufacturer’s instructions. The labeled protein was incubated with test compounds at a final concentration of 200 µM in assay buffer (1x PBS, 0.005% Tween-20, pH 7.4, 2% DMSO) for 15 minutes at room temperature. Samples were then centrifuged to remove bubbles and loaded into 384-well plates for TRIC-based readout. Fluorescence was measured before and after infrared laser activation, and results were expressed as normalized fluorescence (F_norm_).

Selection criteria were applied as previously described^23^. Compounds producing a ΔF_norm_ greater than 10 units, approximately 50% of the CD80 positive control signal, were considered potential hits. Compounds exhibiting autofluorescence, quenching, aggregation, or scan anomalies were excluded. All remaining compounds were classified as hits and used to inform the ML model.

### 3.5. HTS and Hit Selection for the Validation Set Using SPR

HTS of the *Validation Set* was performed using a Biacore™ 8K instrument (Cytiva, Marlborough, MA, USA) as previously described in detail^24^. Briefly, a single-dose binding assay was conducted at 100 µM for 1,056 compounds using a custom method developed in Biacore™ 8K Control Software. Biotinylated human CD28 protein (Cat# CD8-H82E5, Acro Biosystems) was immobilized on a Biotin CAPture Series S sensor chip via Biotin CAPture Reagent (Cat# 28920234, Cytiva). Binding interactions were measured at 25°C in running buffer (1x PBS-P+ with 2% DMSO), with a 60-second contact time and 100-second dissociation phase at a flow rate of 30 µL/min. Solvent correction and anti-CD28 antibody (Cat# K123A, Promega, Madison, WI, USA) were used as controls to ensure data quality and system performance. Data acquisition and analysis were performed using Biacore™ 8K Control and Insight Evaluation Software (Cytiva).

Hit selection was based on kinetic and occupancy-based criteria as previously^24^. Compounds were expected to exhibit rapid association and dissociation behavior typical of small molecules. Those showing minimal or no dissociation during the dissociation phase were flagged as non-dissociating binders (NDB) and excluded. Binding occupancy (LO) was used to further classify responses: compounds with LO >200% were excluded, those between 100–200% were flagged, and those between 50–100% were considered potential hits. Only compounds without non-specific binding (NSB) flags and with LO between 50% and 200% were selected as potential hits and used to validate the ML model.

### 3.6. High-Throughput Screening and Hit Selection for the Evaluation Set Using TRIC Technology

HTS of the *Evaluation Set* was conducted using the Dianthus NT.23 Pico system (NanoTemper Technologies), as previously described in detail and briefly summarized above (*Section 3.3*)^23^.

Selection criteria were applied as previously outlined^44^. Briefly, compounds that produced a ΔF_norm_ greater than three times the standard deviation of the negative control were initially considered potential CD28 binders. After excluding compounds that exhibited autofluorescence, quenching, aggregation, or scan anomalies, the remaining compounds were classified as primary hits. An additional control experiment was conducted for each selected small molecule to ensure no interference with the RED-tris-NTA dye alone - either by quenching its signal, increasing its fluorescence intensity, or altering the TRIC trace. Compounds that exhibited any interaction with the dye alone were excluded from further analysis. The final set of small molecules was then selected for binding affinity measurements and further downstream validation experiments.

### 3.7. MST

Binding affinity measurements were performed using a Monolith X instrument (NanoTemper Technologies, Munich, Germany) employing the spectral shift detection method. Recombinant human CD28 protein (Cat# CD8-H525H3, Acro Biosystems, Newark, DE, USA) was labeled using a His-Tag Labeling Kit RED-tris-NTA 2nd Generation (NanoTemper Technologies, Munich, Germany) according to the manufacturer’s protocol.

Binding experiments were conducted in assay buffer containing 154 mM NaCl, 5.6 mM Na₂HPO₄, 1.05 mM KH₂PO₄, pH 7.4, and 0.005% Tween-20. For binding measurements, labeled CD28 protein was used at a final concentration of 40 nM, and RED-tris-NTA 2nd Generation was added at 20 nM. Test compounds were prepared in DMSO and added to achieve a final DMSO concentration of 1% (v/v). The mixture was incubated for 10 minutes at room temperature prior to measurements.

The Monolith X instrument was configured with optimized parameters for spectral shift detection. All measurements were performed in triplicate using premium capillaries. Initial data processing was performed using the MO. Affinity Analysis software (NanoTemper Technologies), and subsequent data analysis and curve fitting were conducted using GraphPad Prism (GraphPad Software, San Diego, CA, USA).

### 3.8. ELISA

A competitive ELISA was carried out using a commercially available kit from BPS Bioscience (CD28:B7.1[Biotinylated] Inhibitor Screening Assay Kit) to evaluate the ability of test compounds to inhibit the binding of biotinylated B7.1 to immobilized CD28^24^. The experiment was performed following the protocol provided with the kit, and a brief description of the procedure is provided below.

Ninety-six–well plates were coated with CD28 protein at a concentration of 2 µg/mL in PBS (50 µL per well) and incubated overnight at 4°C. The following day, wells were washed with 1× Immuno Buffer and blocked with Blocking Buffer for 1 hour at room temperature to minimize nonspecific binding. Test compounds were added at various concentrations together with biotinylated B7.1 (5 ng/µL), and the mixture was incubated for 1 hour to facilitate competitive binding. Uncoated wells served as negative controls for background signal, while wells containing inhibitor buffer in place of compound served as assay controls. After the binding step, wells were washed thoroughly, followed by the addition of Streptavidin-HRP (diluted 1:1000 in Blocking Buffer) and incubation for 1 hour at room temperature. After a final wash, a chemiluminescent substrate was added, and the luminescence signal was measured immediately using a plate reader in luminescence detection mode.

### 3.9. Molecular docking and dynamics

Molecular docking was performed using Maestro Schrödinger (version 2023.2), with ligand and protein preparation carried out via the LigPrep and Protein Preparation Wizard modules, respectively. The prepared ligands were docked into the previously identified CD28 binding site^24^ using the Ligand Docking Wizard under default parameters. Molecular dynamics (MD) simulations were conducted using Desmond, following the same protocol as described in our previous study^45^. Docking results were visualized using Discovery Studio Visualizer (V.24.1.0), and RMSD plots were generated using Grace software.

## 4. Practical Guidelines and considerations for the Use of HTS-Oracle

HTS-Oracle was developed to enhance compound prioritization in HTS campaigns by improving predictive performance and reducing experimental burden. While the model demonstrated strong performance in both retrospective and prospective validations, its successful implementation depends on careful consideration of several practical and methodological factors.

HTS-Oracle is not a universally applicable model; the model must be retrained for each new biological target using a representative and curated training dataset. Its architecture is target-agnostic, but the predictive utility depends critically on the underlying data. Nonetheless, the pipeline is computationally efficient. For example, inference across a library of 6,000 compounds can be completed in under ten minutes on a workstation equipped with an NVIDIA RTX 4070 Ti GPU and a 32-core CPU, allowing rapid deployment once model training and validation are complete.

To minimize false positives, which can be costly in follow-up experimental assays, HTS-Oracle employs a conservative prediction strategy that prioritizes high precision. This design choice reduces the risk of pursuing inactive compounds but may also increase false negatives, particularly for compounds with borderline activity scores. This trade-off is especially acceptable in low hit-rate settings, such as the CD28 campaign described here (∼1-2%), where experimental validation is resource-intensive and the cost of false positives is high. In contrast, screens with higher baseline hit rates may benefit from adjusted prediction thresholds to maximize recall.

The model architecture is optimized for imbalanced datasets, a common feature of early-stage HTS, and performs well under typical hit-rate conditions (1-5%). In settings with higher hit frequencies, more aggressive post-prediction filtering can yield further reductions in compound volume without sacrificing hit recovery. Conversely, in extremely sparse datasets (<1% hits), predictive confidence may decline, and model performance should be validated rigorously on a case-by-case basis before application to large-scale library screening.

To facilitate community adoption, HTS-Oracle is released as open-source, including both application and training code, along with a pre-trained model available through our GitHub repository. Future work will expand its application to additional target classes and assay types, refine model calibration under varying hit-rate conditions, and explore its integration with orthogonal data modalities such as transcriptomic or structural information. Ongoing updates will aim to support broader reproducibility, adaptability, and sustained model performance across diverse screening contexts.

## 5. Conclusions

In this study, we introduce HTS-Oracle, a retrainable, open-source deep learning–based platform for hit prioritization that integrates multi-modal molecular representations in a unified ensemble framework. By combining transformer-derived chemical language models with traditional cheminformatics features and diverse feature selection strategies, HTS-Oracle offers a robust approach to identifying active compounds while minimizing experimental burden. Its application to the co-stimulatory receptor CD28, a prototypical difficult-to-drug target, demonstrated high predictive accuracy (AUC-ROC = 0.9914, average precision = 0.8565) and a 53-fold enrichment in the top 1% of ranked compounds.

Importantly, HTS-Oracle is designed not merely for retrospective benchmarking but for prospective deployment in real-world HTS pipelines. The model prioritizes precision over recall to reduce false positives and incorporates probabilistic thresholds to support confidence-based decision-making. Its efficient runtime—screening thousands of compounds in under 10 minutes—and straightforward retraining process make it a practical tool for iterative library refinement and target-specific optimization. In prospective screening, HTS-Oracle delivered a hit rate of 8.41%—an ∼8-fold improvement over conventional SPR, TRIC, and mass spectrometry-based HTS platforms—while reducing the number of compounds requiring experimental testing by ∼70%. Two prioritized compounds, Z2770993038 and Z4390463985, inhibited CD28–B7.1 binding with micromolar potency, as confirmed by orthogonal assays. These results are particularly noteworthy given the longstanding challenges in identifying small molecule modulators of CD28, a receptor historically considered intractable due to its shallow and dynamic binding interface.

Beyond this specific application, HTS-Oracle addresses two persistent bottlenecks in drug discovery: the limited success of traditional HTS campaigns and the difficulty of identifying chemically diverse yet potent starting points. By leveraging both contextual and structural features of molecules, the model navigates broad chemical space while maintaining selectivity and interpretability—characteristics essential for translation into medicinal chemistry workflows.

Looking forward, we anticipate that HTS-Oracle will serve as a generalizable blueprint for AI-enhanced virtual screening across a wide range of therapeutic targets. Its open-source availability, modular architecture, and demonstrated experimental validation provide a strong foundation for adoption across academic and industry settings. Future extensions may include integration with protein structural data, bioactivity profiles, transcriptomic signatures, or multitask learning frameworks to further enhance predictive performance and biological relevance.

In conclusion, HTS-Oracle represents a scalable, experimentally grounded, and practically deployable platform that bridges the gap between large-scale virtual screening and focused, resource-efficient hit validation. Its successful application to CD28 highlights the potential of machine learning to redefine early-phase drug discovery—making previously inaccessible targets newly tractable.

## Acknowledgments

This work was supported by the National Institute of Diabetes and Digestive and Kidney Diseases (NIDDK) under grant number R01DK137299.

## Data Availability Statement

All datasets used for modeling are available in our GitHub repository: https://github.com/HTS-Oracle/CD28-project.

## Code availability

The HTS-Oracle Discovery framework, including environment setup, usage tutorial, trained model, system requirements, trained model and the HTS-Oracle application are available as an open-source Python package and via GitHub at: https://github.com/HTS-Oracle/CD28-project.

